# Reconstitution of renal cyst formation in 3D culture reveals a role for AMOT and Yap1 in determining cyst size

**DOI:** 10.1101/2022.03.01.482118

**Authors:** Clark D. Wells, Kevin Lange, Abigail F. Thompson, Wei Min Xu, Sherry G. Clendenon, John S. Underwood, Peter Harris, Britney-Shea Herbert, James Glazier, Angela Wandinger-Ness, Robert L. Bacallao

## Abstract

Despite substantial progress in identifying and understanding causative mutations in autosomal dominant polycystic kidney disease (ADPKD), little is known about subsequent cellular events leading to cyst formation. In prior studies we reported that Cadherin 8, a type II Cadherin, expression is sufficient to induce cyst emergence from HK-2 cells grown as tubule arrays in collagen matrix (1). However, emergent cysts did not exhibit the luminal enlargement observed in ADPKD. In this communication, we reconstitute cyst emergence with consequent cyst lumen expansion in 3D culture by stable co-expression of Cadherin 8 in combination with a constitutively active mutant of YAP1, the key effector of the HIPPO pathway. Specifically, immortalized cells derived from ADPKD cyst epithelia formed cysts with substantially larger lumen sizes when transduced with YAP1-5SA. Conversely, expression of the YAP1 inhibitor, AMOTL1, in these cells resulted in their forming cysts with smaller lumens than control cells. Our data show that cyst formation results from a sequential two-step process consisting of cyst initiation and subsequent cyst expansion. Taken together, cyst initiation induced by Cadherin 8 expression is proposed to result from decreased cell-cell adhesion while cyst expansion is driven by increased Yap1 activity.

## INTRODUCTION

Autosomal dominant polycystic kidney disease (ADPKD) is a relatively common genetic disease associated with kidney enlargement due to the formation of cysts that progressively expand in size until end-stage kidney disease results (2). The mutations responsible for ADPKD cluster within three genes; HmPKD1, HmPKD2 and GANAB (3-5). Of the three loci, 97% of ADPKD cases arise from mutations in HmPKD1 and HmPKD2 (2), which encode polycystin-1 and polycystin-2. Polycystins-1 and 2 form a macromolecular complex in monocilia of renal tubule epithelial cells where they participate in mechanotransduction and exosome signaling to propagate calcium signaling (6, 7). While it is established that a failure of cilia assembly or cilia signaling initiates renal cysts, the subsequent cellular signaling pathway changes that underlie cystogenesis is largely unknown.

Cyst formation results from a localized disruption of morphologic signaling that consequently causes a series of events that disrupt normal tubulogenesis. Cysts initiate as a result of epithelial cells escaping from the planar surface of the tubular epithelia. Such cells fail to complete terminal differentiation and consequently lose both contact inhibition of proliferation as well as apical lumen size control (8, 9). In the classic monograph published by Baert, microdissection analysis of an early stage ADPKD kidney identified two distinct types of emerging cysts (10). Saccular cysts were observed to arise as plaque like colonies of cells from the tubule plane to eventually form a wide base cyst. Stalk-like cysts typically arise from the tubule via a narrow stalk and are connected to a ballooning cell group with a large fluid filled lumen (10). In our prior work we showed that a loss of contact inhibition results in epithelial cells with modestly increased proliferation rates that produce saccular cysts (8, 10). Whereas, virtual modeling predicted that stalk like cysts arise due to decreased cell-cell adhesion between tubular epithelial cells, for instance upon expression of Cadherin 8, a type II cadherin (8).

These concepts were tested in a 3D cell culture model in which the early steps of cyst emergence can be induced. Initially, human renal proximal tubule cells are grown in collagen where they form tubule arrays. Transduction of cells in the tubule arrays with Cadherin 8 using an adenovirus system (1) results in exclusively stalk-like cysts without requiring activation of the endogenous polycystins (1). The effects of Cadherin 8 on cellular cohesion are therefore likely downstream of polycystin1/2 dysfunction. However, cysts produced in this model had uniformly small lumen sizes of less than one micron. This reflected a failure to recapitulate lumen expansion, the primary kidney damaging process in cystogenesis (1). Even though cAMP was previously suggested to promote lumen expansion (11-13), treatment of cysts with compounds such a Br-cAMP or forskolin failed to have an effect. A distinct signaling pathway was therefore predicted to permit or drive lumen expansion.

The Hippo/Yap signaling pathway was originally identified as a master regulator of organ size homeostasis. This pathway was found to link cell-cell adhesion and cellular differentiation cues to the control of cell proliferation rates and survival (14-16). All Hippo signaling converges onto the regulation of the transcriptional co-activators YAP1/TAZ (16, 17). Nuclear entry of YAP1/TAZ is inhibited by their association with AMOT and their phosphorylation by large tumor suppressor kinases (LATs) (14, 18-20). Both YAP1 and AMOT are directly phosphorylated by LATs1/2, which promotes the formation of the AMOT/YAP1 complex with the ubiquitin ligase AIP4. Ubiquitination of YAP1 results in its cytosolic sequestration as well as targeting for degradation (21). Because AMOT also directly binds to members of the PAR apical polarity complex, it coordinates YAP1 regulation with the control of apical specification. This is due in large part by targeting both apical polarity and HIPPO signaling factors to recycling endosomes versus the tight junction (22). Because AMOT coordinates polarity and HIPPO signaling, it was predicted to have a role in tubule lumen size control. In this report we examine the potential of YAP1 and AMOTL1 protein expression to control the morphologic phenotype of cyst formation in 3D culture. Heterologous expression of a constitutively active YAP1 mutant was found to be sufficient for cells to reconstitute cyst enlargement when grown in a 3D matrix. These results support a model where cyst emergence occurs in response to alterations in cadherin expression and is followed by a second step wherein dysregulated Yap1 activity drives cyst lumen size growth. Furthermore, AMOT expression was shown to counter the effects of YAP1 activity and to decrease cyst size in immortalized ADPKD cells grown in 3D matrix.

## RESULTS

The importance of cadherin expression in tissue morphogenesis determination (23) led us to evaluate cadherin expression in immortalized cyst derived human ADPKD epithelia. Studies found that cadherin 8 was aberrantly expressed in cyst epithelia and *in situ* in cysts examined from pathological sections of end-stage ADPKD kidneys (1). However, dysregulated Cadherin 8 expression could be the result of adaptations to cell culture conditions or from trans-differentiation in tissues from adaptations to uremia (24-26). To revisit this question, Cadherin 8 expression was examined in kidney sections from the murine Pkd1RC/RC model of renal cystic disease. The PKD1RC/RC model is characterized by a slowly progressive form of polycystic kidney disease (27). Staining of kidney sections obtained from post-natal day one mice with anti-cadherin 8 antibody showed specific cadherin 8 expression in cysts versus antibody alone controls (Figure 1A, B). The early presence of cadherin 8 in cyst epithelia underscores a potential role for cadherins in cyst morphogenesis.

**Figure 1:**
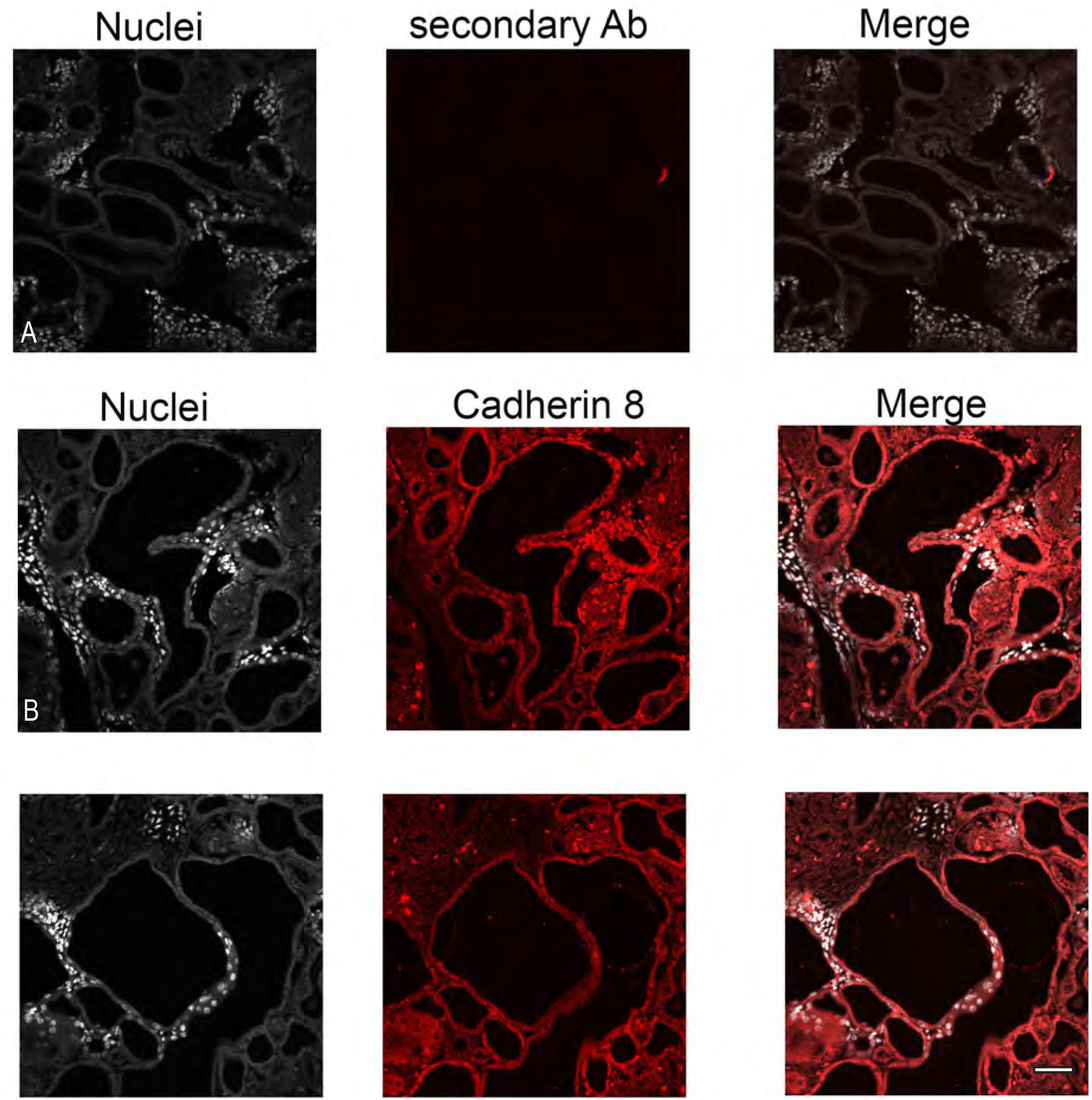
Cadherin 8 is expressed in the cysts of PKD1RC/RC mouse kidneys. Kidney sections made from post-natal day 1 pups are stained with antibodies that bind murine cadherin 8. All kidney sections are labeled with Hoechst 33342 (grey). Figure 1A: Kidney section label with Texas red secondary antibodies only. Figure 2B: Kidney sections labeled with anti-cadherin 8 and nuclear staining. Cystic lumens (c) are labeled, and arrows show cadherin 8 staining in cyst lining epithelial cells. Bar=10 microns.

The effects of changes in cell-cell adhesion on cyst formation were spatially modelled using the CompuCell3D software package. Overall, cyst emergence and subsequent cyst morphology was indicated to be predicated on a loss of contact inhibition of cell proliferation and decreased cell-cell adhesion (8). Analysis of signaling pathways that transmit cell adhesion state on to the regulation of cell proliferation pointed to the Hippo/Yap signaling cascade (Figure 2A). Potential dysregulation of HIPPO signaling proteins during cyst expansion was probed by comparing the levels of proteins in this pathway between normal human kidney epithelial cells (NK), human ADPKD non-cystic cells (NC-ADPKD) and patient matched ADPKD cystic cells (C-ADKPKD) (28). Immunoblot analysis shows reduced levels of SAV1, MST2, LATS1, YAP1, and P-YAP (Ser 127) in C-ADPKD cells relative to normal kidney NK cells (Figure 2B). In contrast, levels of P-MOB1 (Thr 35) and P-YAP (Ser 397) were increased in C-ADPKD cells as compared to NK cells (Figure 2B). Whereas the levels of SAV1, MST1, MOB1, and P-MOB1 (Thr35) were relatively similar between NK and NC-ADPKD cells (Figure 2B). Alternatively, MST2, LATS1, YAP1, P-YAP1 (Ser 127) were lower in NC-ADPKD cells versus NK cells (Figure 1B). While our data points to a loss of HIPPO kinases in cyst derived ADPKD cells, total levels of YAP/TAZ were also lower in cystic epithelial cells. This observation may be explained by the increased levels of P-YAP (Ser 397), which indicates that YAP1 is being targeted for degradation (18, 29). Upon examination of the complex interactions between components of the Hippo/Yap pathway as shown in Figure 2A, became apparent that computer modeling of the numerous potential changes in the interactive pathway would be challenging, so we explored the functional effects of Yap1 on lumen size control in our cell culture model systems.

**Figure 2:**
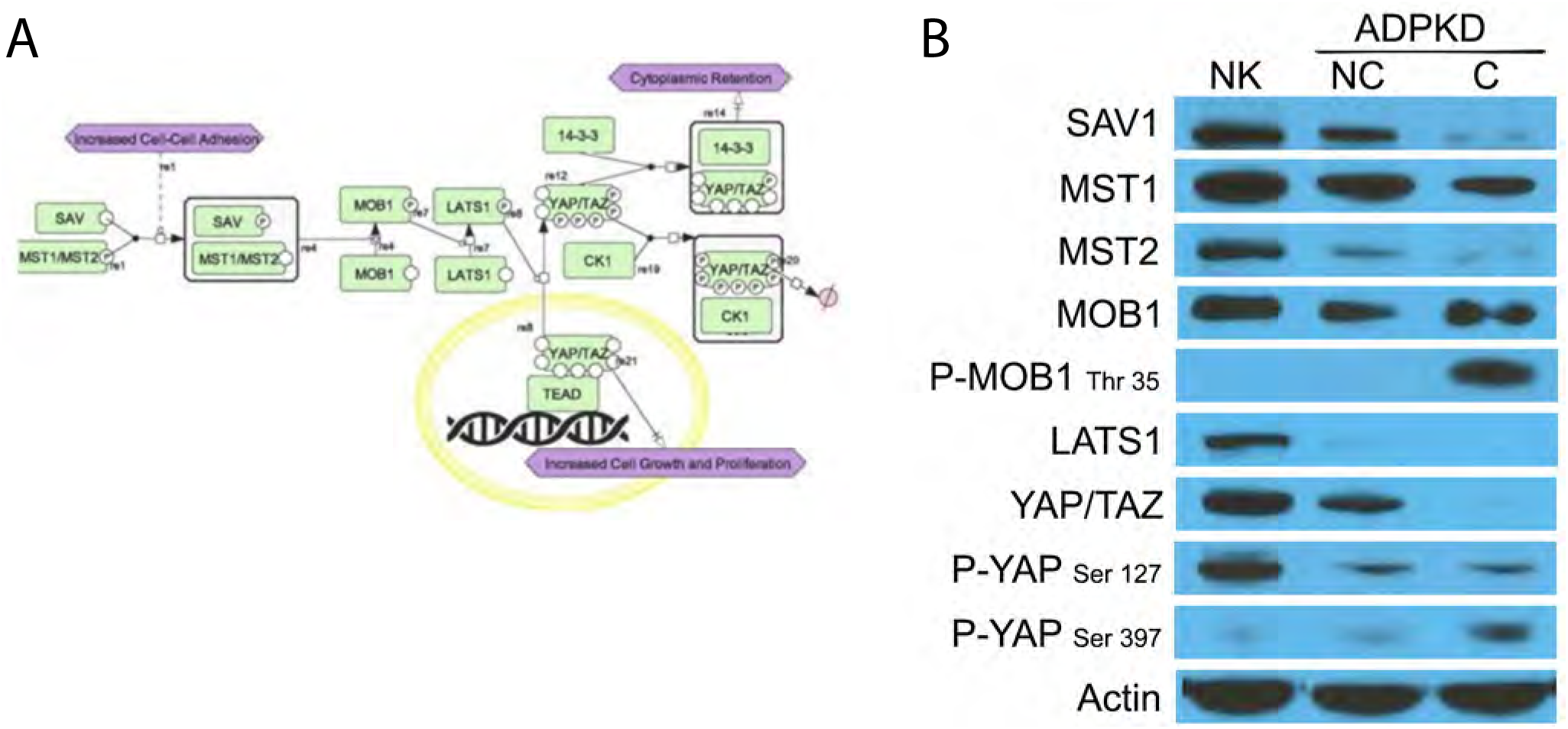
HIPPO/YAP signaling system and immune blot analysis of HIPPO/YAP signaling protein expression in human renal cell lines. A: Cell designer schematic of Hippo/Yap signaling. The regulatory components of the pathway are shown, and this schematic can be used as a modeling system for predictive outcomes in the signaling system. B: Immune blot analysis of Hippo signaling protein expression in normal human proximal tubule cells (NK), non-cystic ADPKD cells derived from normal appearing renal tubules (NC), and cystic epithelial cells derived from the same ADPKD kidney (C). Antibodies to Sav1, MST1, MST2, MOB1, phospho-MOB1 (Thr35), LATS1, YAP/TAZ, phospho-YAP (Ser127), and phosphor-YAP (Ser 397). Actin loading controls are shown in the bottom panel.

AMOT or angiomotin, is an adaptor protein that shuttles between the tight junction and a juxta-nuclear compartment where it controls apical membrane trafficking of Patj, Pals and Par3 polarity complex proteins (22, 30). AMOT is also complexed with YAP at the tight junction where YAP plays a role in contact inhibition (31, 32). We sought to test the possibility that YAP plays a heretofore unrecognized role in apical lumen size control. To test this hypothesis, we utilized our 3D cell culture model where exogenous cadherin 8 expression initiates cyst emergence from pre-formed tubule arrays (1). The model recapitulates cyst initiation from loss of contact inhibition but suffers from a failure to produce cysts that have large lumens, a critical clinical feature of ADPKD. The model employs HK-2 cells grown in tubule arrays in porcine tendon collagen and cyst emergence is activated by adenovirus driven human cadherin 8 expression. We transduced HK-2 cells with lentivirus containing constitutively active YAP1-5SA (20, 33), and grew the cells in collagen matrix. We found that YAP1-5SA transduced HK-2 cells form tubule arrays indistinguishable from tubule arrays formed by normal HK-2 cells (Figure 3A). However, when cyst emergence is initiated by cadherin 8, cysts emerge from the tubule but lack resolvable lumens especially in when one examines the YZ and XZ planes (Figure 3 B). In cultures with HK-2 cells transduced with Yap1 5AS and then cyst emergence is induced with cadherin 8, cysts form with large lumen spaces (Figure 3 C). In contrast, YAP1-5SA transduced cells expressing additional N-cadherin driven by adenovirus infection, fail to form cysts (figure 3D). Furthermore, HK-2 cells transduced with cyan fluorescent protein and then infected with cadherin 8 adenovirus, failed to form emergent cysts (supplemental figure 1). This result shows that the fully realized cyst emergence with enlarged lumens is specifically determined by a two-step event requiring Yap1 and cadherin 8 activity and is not a result of a cellular response to an exogenous gene expression event. This finding further supports our conclusion that cyst lumen expansion is dependent upon YAP1-5SA.

**Figure 3:**
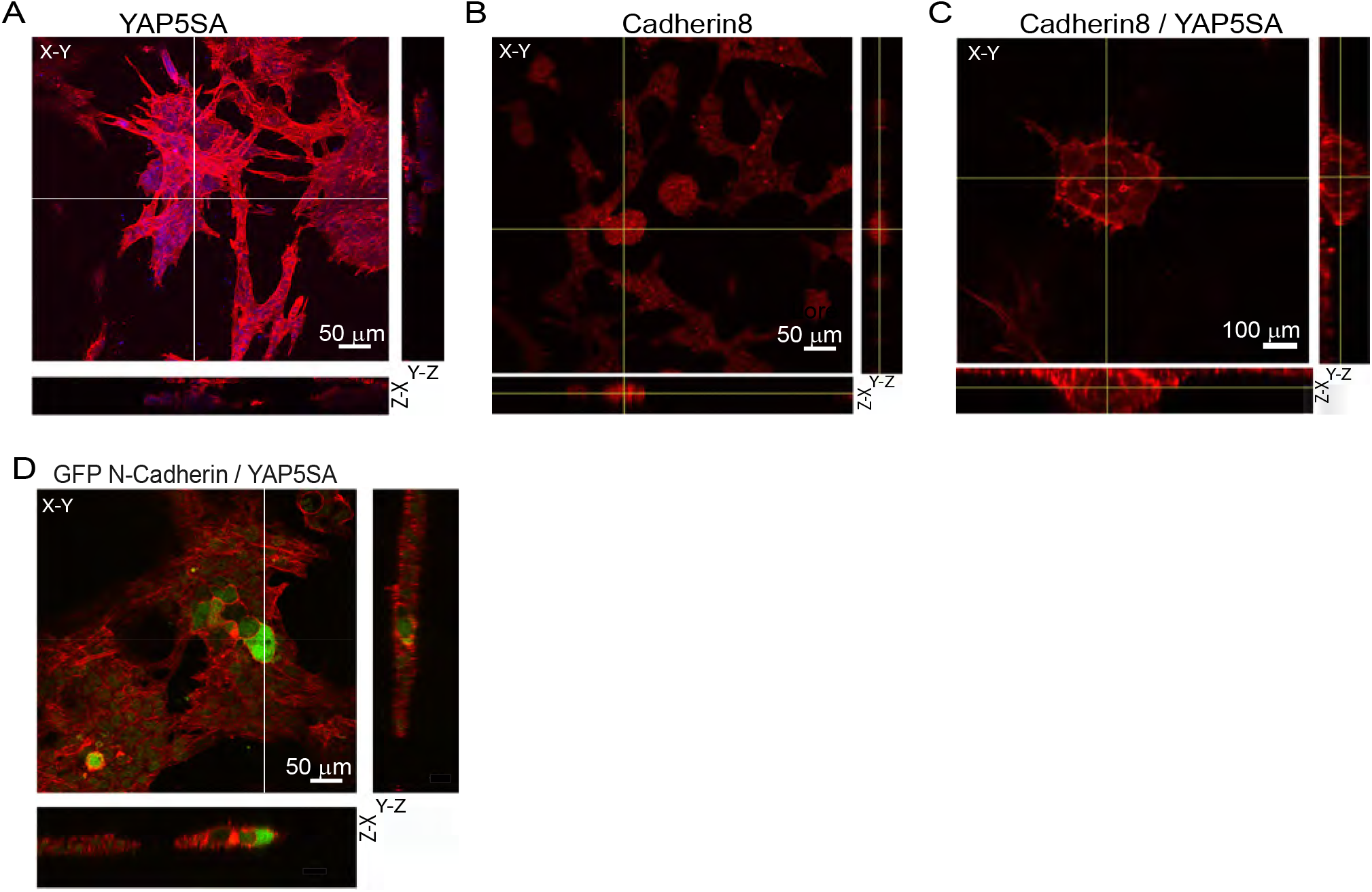
HK-2 cells transduced with YAP1-5SA, grown in 3D culture and expressing cadherin 8 form cysts with enlarged lumen domains. Cells labeled with rhodamine phalloidin (red). Vertical lines in the X-Y images are the section site for the Y-Z orthogonal images. Horizontal lines in the X-Y images are the section site for the X-Z orthogonal images. A: HK-2 cells transduced with YAP1-5SA without cadherin 8 expression form tubule arrays in 3D culture. Orthogonal images in the XY, XZ and YZ planes. Bar=50 µm. B: HK-2 cells forming cysts following cadherin 8 expressing adenovirus injection. No cyst lumen is apparent in the orthogonal sections in the YZ or XZ planes. Bar= 50 µm. C: HK-2 cells transduced with YAP1-5SA with subsequent cadherin 8 adenovirus infection form a cyst with a single lumen. Bar= 100 µm. D: HK-2 cells transduced with YAP1-5SA with subsequent N-cadherin-GFP adenovirus infection fails to form a cyst. Bar=50 µm.

In prior studies we showed that PKD Q4004X cells form cysts when grown in Matrigel (28). While this cell line does not express cadherin 8 it has lower cadherin 1 expression levels as compared to normal age and sex matched immortalized kidney cells (NHPTK) (28).To evaluate the effect of YAP in a human cyst epithelial cell line, PKD Q4004X cells were transduced with YAP1 5SA and grown in Matrigel for 7 days (28). YAP1-5SA transduced cells formed large cysts in culture as compared to mock transduced cells (Figure 4A, right panel versus left panel). The mean calculated area of cysts from the YAP1-5SA transduced PKD 4004X cells was 6,619.66 ± 773.78 µm2 as compared to 177.89 ± 7.16 µm2 (p value <0.01) (Figure 4B).

**Figure 4:**
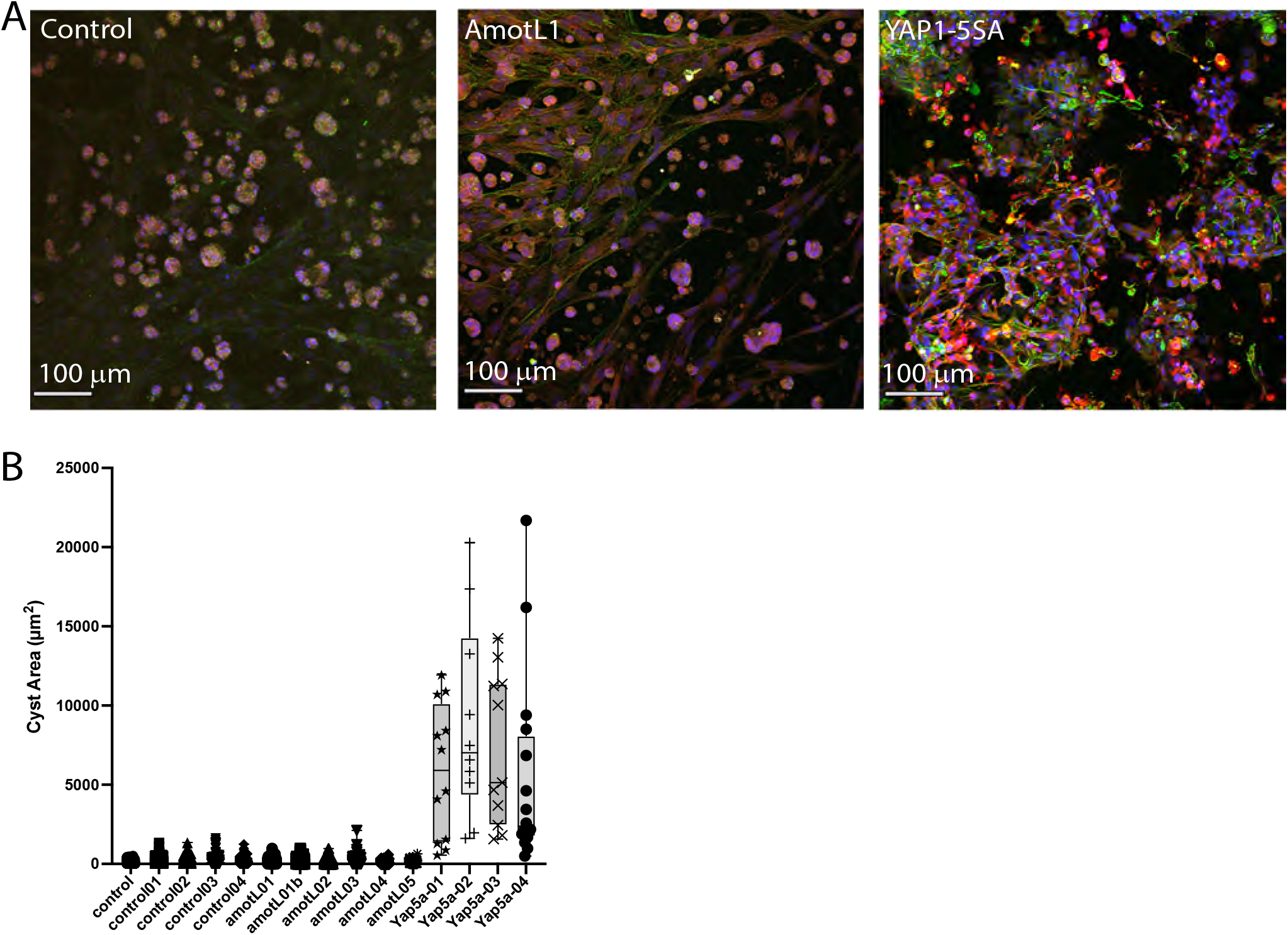
Cyst size in PKD Q4004X cells is increased by YAP1-5SA or decreased by AMOTL1 transduction. PKD Q4004X cells grown in Matrigel for 7 days. Cells are either transduced with vehicle, AMOTL1 or YAP1-5SA. A: Control untransduced cells. B: PKD Q4004X cells transduced with AMOTL1. C: PKD Q4004X cells transduced with YAP1-5SA. Green channel is actin, blue channel nuclei Bar= 100 µm.

The interaction of AMOT with YAP is a control node in cytosolic YAP function (34, 35). We reasoned that expression of constitutively activated AMOT would inhibit lumen size expansion in ADPKD cells (22, 36). AMOT expression in NHPTK and PKDQ4004X cells was equivalent (Supplemental data figure 2A), so to test the effect of AMOT on lumen size, PKDQ4 004X cells were transduced with AMOTL1 (S262E). PKD Q4004X cells transduced with AMOTL1 (S262E) and grown in Matrigel for 7 days formed cysts with an average area of 122.77 ± 6.0 µm2 which is significantly less than the area observed the parental cell line (177.89 ± 7.16 µm2) (p value <0.01) (Figure 4B and Supplemental Figure 2B).

## DISCUSSION

In this study we use an immortalized human proximal tubule cell line derived from phenotypically normal kidneys to reconstitute features of early cyst formation first delineated by Beart and Steg (37). Cellular reconstitution systems have been used in the past to delineate features of pathways in simpler cellular systems. For example, fibroblast cells expressing villin develop features of microvilli (38, 39). We take a similar approach in this communication to understand the early steps of cyst formation from cysts emerging from a preformed tubule array to the enlarging cyst lumen. It is the latter feature of our model system that recapitulates the salient feature of ADPKD; enlarging cysts with a concomitant increase in lumen size (4). In our studies we show that transduced HK-2 cells with YAP1-5SA form tubule arrays indistinguishable from untransduced cells when grown in 3D culture. Subsequent infection with adenovirus that creates focal cadherin 8 expression is necessary to initiate cyst emergence. This result agrees with our prior published result, where cadherin 8 expression is sufficient to initiate cyst emergence (1). However, while cadherin 8 can initiate cyst emergence, the resultant cysts lack enlarged lumens. It is the combination of YAP1-5SA and cadherin 8 expression that results in cysts with enlarged lumenal spaces (Figure 3). Our results identify two steps that are necessary for cyst formation; altered cell adhesion likely initiated by an aberrant cadherin that propels cyst emergence followed by loss of normal apical lumen size control (1, 8). In our examination of cadherin expression in cyst epithelia, cadherin 8 is not the only atypical cadherin expressed, however we have found it expressed in end-stage ADPKD kidney cysts and in the murine *PKD1RC/RC* kidneys from post-natal day one mouse kidneys (Figure 1). Cadherin 8 expression in an early stage PKD1 model, coupled with our results suggests that cadherin 8 is a morphogenic inducer of cystogenesis (1).

### Lumen size control in biological systems

Human autosomal dominant polycystic kidney disease is characterized by enlarging cystic structures within the kidney parenchyma (4). Cyst enlargement has been linked to a secretory phenotype manifest by the cyst epithelia lining the cyst lumen (11, 40). Strikingly the normal nephrons in autosomal dominant polycystic kidneys have phenotypically normal appearing lumen spaces, this suggests that cystogenesis results in a loss of lumen size control.

Lumen size in organs with regions surrounded by a layer of epithelial or endothelial cells is determined by a complex interplay between extracellular matrix, cell proliferation controls, planar polarity and apical-basal polarity elements (41). In 3D culture studies employing Madin-Darby canine kidney cells, apical polarity elements such as crb3, PALs, PATj have been found to determine lumen size and or lumen number in cell growths (41). These apical determining proteins are found in larger complexes in apical recycling endosomes in which AMOTL1 has been shown to play a critical adaptor role, targeting these endosomal complexes to the apical-basal border demarked by the tight junction (22). Apical lumen size is negatively regulated by Scribble and very likely elements that control apical membrane recycling (41). It has also been shown to affect cyst size due to its inhibitory role in the Hippo/YAP pathway (42). Our results highlight a role for the relative amounts of AMOTL1 and Yap in cyst size in our model system.

Another feature of cyst enlargement is coordinated cell proliferation. In our computer model studies, we found that cell proliferation must be in a constrained rate for cyst formation to occur. Our parameter sweep studies pointed to a decrease of doubling time by 30% relative to normal was the optimum change in proliferation for cystogenesis (8). Among the candidate signaling systems that may drive cyst growth, the Hippo/Yap pathway is an attractive candidate for evaluation since it integrates cell-cell adhesion, contact inhibition, and cell proliferation in a relatively controlled way (15, 31, 43). Notably, Yap has been identified to transcriptionally regulate c-myc in a polarity axis that modulates renal cyst formation in a relevant model of murine *pkd1-/-*(27). Additional elements of the Hippo-Yap pathway have also been shown to play a critical role in cystogenesis, the PDZ binding motif transcriptional activator (TAZ) binds to β-catenin with polycystin-1. In polycystin-1 deficient cells TAZ binds to AXIN-1 and thereby increases β-catenin transcriptional activation of c-myc, a process that TAZ also directly stimulates transcriptionally.

Our results directly link a duality of roles for the Hippo/Yap pathway in cystogenesis. Elements of the pathway such as TAZ and YAP are activating cell proliferation in a relatively controlled manner that is distinct from a full-blown malignant transformation (44). Our findings also point to a direct role in apical lumen size control vial modulation of apical polarity complexes that reside in the apical recycling compartment whose rate of apical delivery is controlled by AMOT (22). This suggests that therapies targeted to modulating AMOT or YAP are likely to impact both cell cyst proliferation and cyst lumen size.

## METHODS

Immune-blot analyses were carried out on lysates made from confluent PKD Q4004X and HmPTK cells using antibodies for SAV1, MST1/2, MOB1, p-MOB1 (thr 35), LATS1, YAP/TAZ, p-YAP (ser 127) and p-YAP (ser 397). HK-2 cells transduced with lentivirus bearing constitutively active YAP5SA were grown in 3D culture as previously described (1). Cyst emergence was activated by expression of cadherin 8 upon transduction of adenovirus bearing Cahderin-8 (1). PKD Q4004X cells were transduced with lentivirus expressing either YAP5SA or the constitutively active AMOTL1 (S262E) mutant and grown in Matrigel for 14 days (28). Cyst size was assessed by confocal microscopy and image processing with Image J (45).

### Antibodies, chemical and molecular biology reagents

All fluorescence conjugated secondary antibodies were obtained from Jackson ImmunoResearch (Warrington, MA). Anti-MST-1, 2, SAV1, MOB1, LATS1, YAP/TAZ, phospho-MOB1 (Thr35), phospho-YAP (Ser 127), phospho-YAP (Ser 397) were purchased from Cell Signaling Technology (Danvers, MA). Rhodamine conjugated phalloidin and Hoechst 33342 were purchased from ThermoFisher Scientific (Waltham, MA). Cell matrix was obtained from the Nittan Gelatin Company (FUJI-Wako, Richmond, VA). Matrigel was supplied by ThermoFisher Scientific (Waltham, MA).

### Cell lines and culture conditions

HK-2 and 293T cells were purchased from ATCC and grown in continuous culture as previously described (1, 35). PKD Q4004X and NHPTK cells were maintained in culture as previously described (28). NC-PKD and C-PKD cells are telomerase immortalized cell lines derived from an early, chronic kidney disease stage I, polycystic kidney with a heterozygous truncation mutation at Q2556X of the HmPKD1 gene (46). NC-PKD cells were isolated from normal tubules dissected from the kidney and C-PKD cells were isolated from cysts of ADPKD kidneys (46). The cells were maintained in culture under the same culture conditions as PKD Q4004X and NHPTK cells (28). All cell lines have been validated by IDEXX laboratories and are mycoplasma free. 3D cell cultures were produced as previously described for Matrigel in the case of PKD Q4004X cells (28) or HK-2 cells (1).

### Immunoblots

Antibodies to Yap/TAZ, MST1, MST2, LATS1, phospho-Yap (Ser297), phospho-MOB1(Thr35), MOB1, SAV1, phospho-Yap (Ser 127) were obtained from Cell Signaling Technology (Danvers, MA). Cell lysates were made from confluent cultures of NHPTK, NC-ADPKD and C-APDKD cells using RIPA buffer (140 mM NaCl, 10 mM Tris-Cl; pH 8.0, 1 mM EDTA,0.5 mM EGTA, 0.1% sodium dodecylsulfate, 1% Triton X-100, 0.1% sodium deoxycholate) supplemented with 0.5 mM AEBSF and 10 ug/ml chymostatin, leupeptin, pepstatin and aprotinin. Protein concentration of each lysate was measured using the BCA protein assay kit according to manufacturer’s directions (Thermo-Fisher Scientific, Waltham, MA). Proteins were separated on a 10% SDS-PAGE gel with equivalent amounts of protein loaded in each lane (Thermo-Fisher Scientific, Waltham, MA). After transfer to nitrocellulose, blots were blocked with 1% newborn calf serum in 1% Triton X-100 dissolved in tris buffered saline. All primary antibody incubations were performed with antibodies dissolved in 1% Triton-X 100, 1% newborn calf serum and tris-buffered saline. Horse radish peroxidase conjugated secondary antibodies were purchased from Jackson ImmunoResearch Laboratories (West Grove, PA). Blots were developed with Super Signal according to manufacturer’s directions (Thermo Fisher Scientific, Waltham, MA). To assess equivalence of protein loading, after blots were developed, they were stripped with ReBlot Plus (Millipore Sigma, St. Louis, MO). Blots were then probed with anti-actin and developed as described above.

### Lentivirus Transduction

Lentivirus constructs containing Yap5A and AMOTL1 were made as previously described (35). Viral transduction was performed by adding culture supernatant collected from transfected 293T cells, adding polybrene to the culture supernatant and then added to 80% confluent target cells with a multiplicity of infection of 10:1. Studies were conducted in transduced cells that were passaged no more than twice following transduction.

### Image Acquisition

3D cell cultures were washed with phosphate buffered saline and then fixed with 4% paraformaldehyde dissolved in phosphate buffered saline, pH 7.,4 for 30 minutes. Fixation reactions were quenched with 100 mM NH4Cl dissolved on phosphate buffered saline. Samples were labeled with primary and secondary antibodies dissolved in 140 mM NaCl, 10 mM Tris-Cl; pH 8.0, 0.5% Triton X-100 (TBST) and 0.2% fish skin gelatin (Sigma Aldrich, St. Louis, MO). After incubating with primary or secondary antibodies the samples would be washed 6 times with TBST, with 30-minute wash incubations performed at room temperature. Hoechst 33342 and rhodamine phalloidin was added to all secondary antibody solutions. Following labeling, all samples were fixed with 2% paraformaldehyde dissolved in phosphate buffered saline and mounted in 50% glycerol/phosphate buffer saline with 1 mg/ml 1,4-diazabicyclo [2.2.2] octane.

Images were collected with an Olympus Flowview FV1000 confocal module mounted on an Olympus iX 81 inverted microscope (Olympus Life Sciences, Center Valley, PA). Excitation light was provided by a Spectraphysics DeepSee tunable titanium-sapphire laser at 800 nM femtosecond pulsed laser (Spectra-Physics, Santa Clara, CA). All images were collected with a 20X water immersion lens (N.A 0.95, working distance= 1.0 mm) (Olympus Life Sciences, Center Valley, PA). Data collection from all samples was performed under identical intensity, black level and line scan settings. Image processing was performed using FIJI open-source software running on an HP laptop (45).

### Analysis of cyst areas

Cyst areas were determined by collecting 3D volumes of PKD Q4004X cells grown in Matrigel for seven days. Image stacks were collected to match the z distance with x-y distance per pixel to obtain cubic voxels for analysis (47). The rolling ball background subtraction was used to remove spatial variation in background intensities and cyst regions were identified using Weka Trainable Segmentation (48). Using the Analyze Particles command, cyst region areas were measured, and data was stored for each cyst area in the region of interest manager. The cyst area data from the ROI manager was then saved as a csv for statistical analysis.

## Acknowledgements

RLB, CRW conceived and performed experiments, interpreted results and edited the manuscript. BSH, AWN created the immortalized cell lines. PH provided murine samples, interpreted results, and edited the manuscript. JSU, AT, WX and KL performed experiments. SC and JC analyzed computer simulations. SC planned experiments and performed image analysis. All image acquisition was performed at the Indiana Center for Biological Microscopy which is supported by an O’Brien Award from NIDDK (P30 DK079312). RLB is supported by awards from Dialysis Central International and VA Merit, I01BX005184. AWN is supported by NCI P30 CA118100 and immortalized cells were generated under previous NIDDK R01DK50141. CDW is supported by R01GM137656. The views expressed in this article are those of the authors and do not necessarily reflect the position or policy of the Department of Veterans Affairs or the United States government.

---

Supplemental Figure 1: Cyan fluorescent protein (CFP) transfected HK-2 cells grown in 3D culture and infected with cadherin 8 expressing adenovirus do not form cysts. Cyan fluorescent protein transfected HK-2 cells were plated in pig tendon collagen (FUJI Film Wako Chemicals, USA,). Cyan fluorescent protein positive regions of the culture were identified and cadherin 8 expressing adenovirus. X-Y plane of (CFP and nuclear staining (blue channel) and actin (red channel).and orthogonal planes (Y-Z) and (X-Z) plane are shown also. Bar=50 µm.

Supplemental Figure 2: A. AmotL1 expression in PKD Q4004X and NHPTK cells. NHPTK and PKD Q4004X cells were grown in 3D collagen matrix for 7 days. Matrix was digested with dispase, and extracts were made with hot sample buffer. After SDS_PAGE and transfer to nitrocellulose, the blots were probed with anti-AMOTL1. Lanes 1-3: NHKPTK lysates, lanes 4-6: PKD 4004X lysates. Lower molecular weight bands are proteolytic fragments from the matrix digestion. B: 3D projections of PKD Q4004X cells grown in 3D culture. Top panel: PKD Q4004X cells, untransfected control, Middle panel: PKD Q4004X cells transfected with AmotL1, Lower panel: PKDQ4004X transfected with YAP5AS.

