## Supplementary figures and images for "Reconstitution of renal cyst formation in 3D culture reveals a role for AMOT and Yap1 in determining cyst size"

### Supplemental Figure 1

Supplementary Figure 1

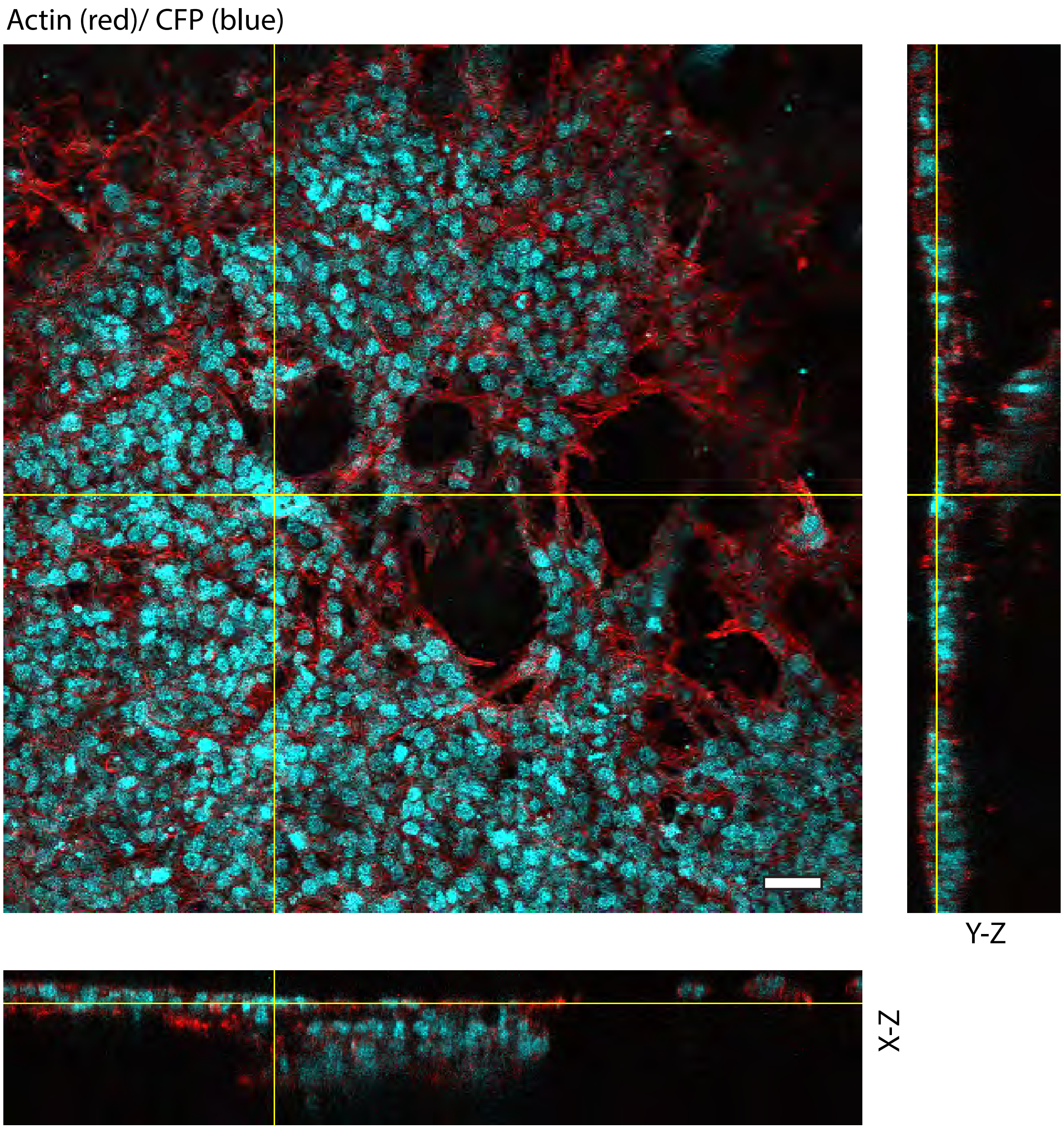

### Supplemental Figure 2

Supplementary Figure 2

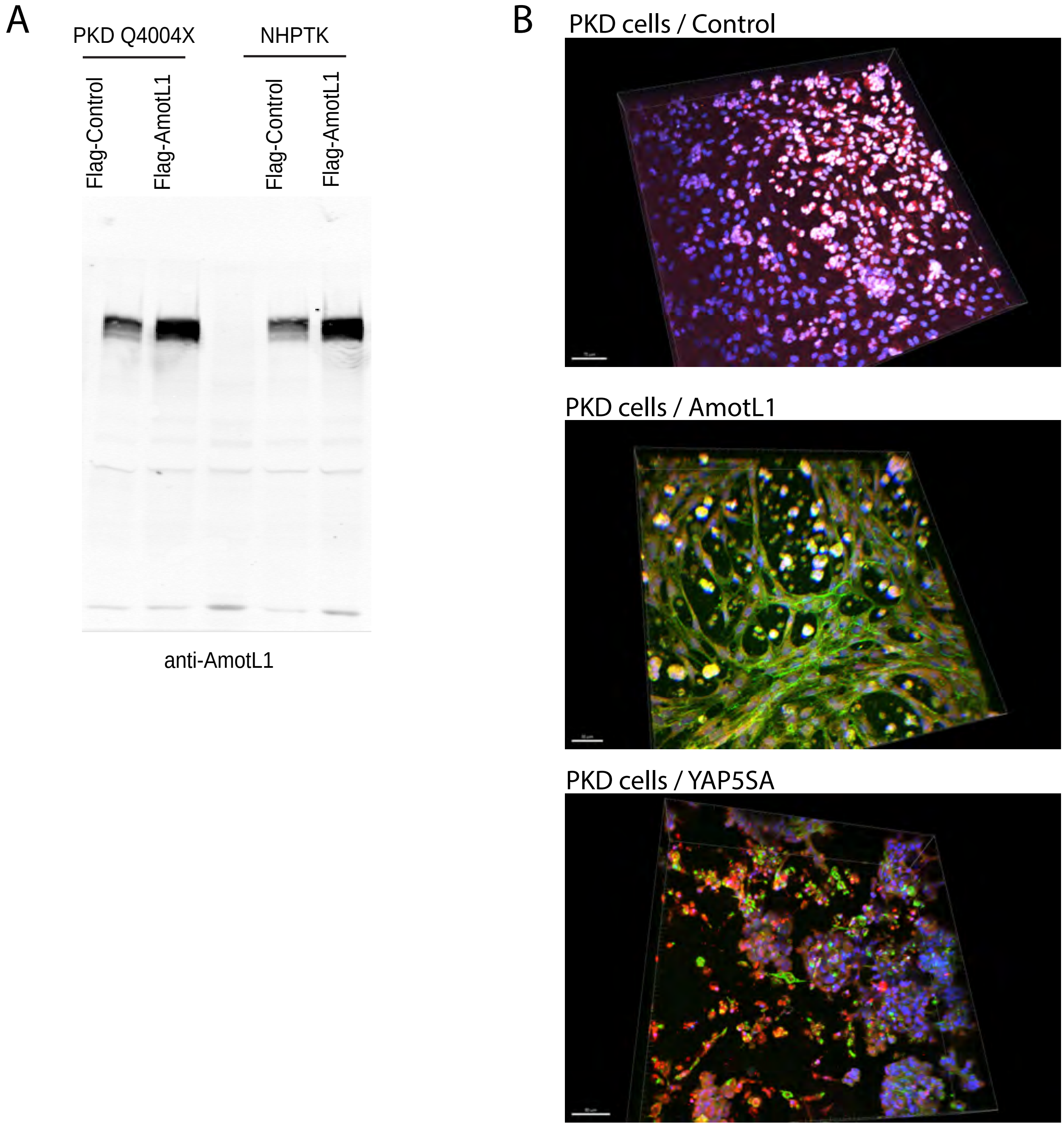
